# Apparent failure and cryptic success of disease control via intermediate host population reduction

**DOI:** 10.64898/2025.12.08.693001

**Authors:** Nadia Raytselis, Maya Risin, Layla Steinbock, Logan T. Story, Jane Miller, Ta’Nyia Heard, Kevin Xie, Ella Rintala, Inesha Gupta, Stephanie Gutierrez, Alexander Strauss, David J. Civitello

**Affiliations:** Department of Biology, Emory University, Atlanta, GA, 30322, USA; Odum School of Ecology, University of Georgia, Athens, GA, 30602, USA; Center for the Ecology of Infectious Diseases, University of Georgia, Athens, GA, 30602; River Basin Center, University of Georgia, Athens, GA, 30602

**Keywords:** population dynamics, negative density dependence, pest management, compensation, overcompensation, disease control

## Abstract

All populations experience density dependence, lest they grow infinitely. However, elucidating what forms of density dependence act most strongly for a given species remains incredibly challenging. Identifying the mechanisms that regulate population density is particularly relevant to pest control, which often causes high mortality, reducing population density but enabling recovery when survivors are freed from intraspecific competition and density dependent vital rates (e.g., birth, death, and maturation rates). Understanding these demographic responses is critical for effective pest management, especially when specific life stages of pests differ in the harm they cause. Guinea worm disease (GWD) is a neglected tropical disease with an obligate copepod intermediate host, and importantly, only large-bodied copepods can be infected with GW parasites. GW disease has been the target of control for several decades, with main management strategies including the use of a chemical larvicide, Abate™, which is applied to water bodies to cull copepod populations. Despite the wide application of Abate in GW endemic countries, little is known about long-term copepod population dynamics in response to Abate™. Thus, we evaluated this mortality-based management of intermediate copepod hosts to control GW transmission, combining mathematical models with a population dynamics experiment designed to mimic Abate™ pesticide control methods across a range of intensities. Despite initial reductions in both total and stage specific population densities, copepod populations recovered so rapidly that control appeared to fail, enabled by extremely fast maturation rates in low-density populations. Much to our surprise, model simulations of GW transmission showed that although total and stage-specific densities rebounded, infectious adult copepods —the proximate cause of infections—were strongly suppressed, indicating cryptic success. This effect was enabled by the 15-day developmental period of GW larvae within copepods, which blocks the accumulation of infectious copepods between interventions. Ultimately, this study highlights the importance of understanding mechanisms of density dependence when designing and optimizing pest control interventions, as well as interpreting counterintuitive consequences of interventions.

## Introduction

Elucidating the mechanisms that regulate populations is a fundamental goal of population ecology. Negative feedbacks on population growth emerge because of competition for space (Sebens 1982) and resources (Svanbäck and Bolnick 2007), degradation of the environment (Calizza et al. 2017), or other mechanisms that reduce individual performance as density rises (Lu and Wang 2025, Stockseth et al. 2025). Reductions in birth rates (Gotelli 2008) and increases in death rates are ubiquitous inhibitors of population growth (Hixon and Jones 2005). In populations with stage structure, negative density dependence can also arise through maturation rates or cannibalism (Park et al. 1965, Dennis et al. 2001, Walker et al. 2021). Despite the reality that all populations experience negative density dependence (lest they increase infinitely), it remains challenging to identify which form(s) operate most strongly for any species and few studies attempt to do so (Kaspersson et al. 2013).

The forms of density dependence are particularly relevant for pest control, because interventions are designed to reduce pest density. This reduction is often followed by rapid recovery, driven by relaxed density dependence (Karatayev et al. 2015) or potential acquisition of resistance (Cao et al. 2025), limiting benefits and necessitating repeated interventions (McIntire and Juliano 2018). The ensuing dynamics of repeated pest suppression and recovery can be highly sensitive to density-dependent vital rates (Smith 1997). For example, strong negative density dependence can cause compensation or overcompensation, the phenomena in which control efforts actually increase the density or biomass of some or all life stages (Karatayev et al. 2015, de Roos 2020, Grosholz et al. 2021). Even when interventions successfully reduce density, populations can rebound faster than expected if the remaining individuals are released from density-dependent reductions in maturation or birth rates (Karatayev et al. 2015). Such responses may be strongest in stage-structured populations, where removal of more competitive classes, or reductions in density overall, elicit stage-specific compensatory responses (Karatayev et al. 2015). Thus, delineating the form(s) and magnitude of density dependence can improve management of the ecological issues that pests cause, such as disease transmission, crop losses, and biodiversity harm.

Here we combine an experiment manipulating pest control intensity with mechanistic modeling to identify which density-regulating mechanisms affect copepods, the intermediate hosts of the Guinea worm (GW) parasite, *Dracunculus medinensis*. Copepods exhibit stage-structured populations (van den Bosch and Gabriel 1994) and experience repeated control interventions due to their role in GW transmission (Gindola et al. 2022). Guinea worm disease (GWD) is a globally impactful infectious disease, causing disability and negative economic impacts in endemic areas (Pellegrino et al. 2022). Cyclopoid copepods are obligate intermediate hosts that become infected through consumption of GW larvae and facilitate transmission to humans and other definitive mammalian hosts (Simonetti et al. 2023). GWD has been targeted for eradication for several decades (World Health Organization 1986), with few cases remaining across five endemic countries (World Health Organization 2025). In GW-endemic regions, application of the pesticide Abate™ (active compound: temephos) is a major component of eradication efforts (Gindola et al. 2022). Abate™ can cause >99% immobilization of copepods in laboratory studies (Grunert et al. 2022), although copepod species vary widely in their sensitivity (Hanazato et al. 1989, Leboulanger et al. 2011). In Abate^TM^-treated sites, copepod populations often exhibit immediate reductions in density, but routinely recover, necessitating application at 28-day intervals to support eradication (Gindola et al. 2022). The density-dependent demographic responses of these intermediate hosts – along with implications for disease control – are unknown.

Copepod stage structure has implications for both their population dynamics and GW transmission. Neonatal copepods (nauplii) are too small to consume GW larvae, but larger juveniles (copepodites) and adults can consume parasites and become infected (McCullough 1983, Bimi 2007). Once ingested, parasites molt twice over 10-16 days within the copepod before maturing into third stage larvae, the only stage infectious to definitive hosts that imbibe infected copepods (Gonzalez Engelhard et al. 2021). Additionally, adults can cannibalize nauplii (van den Bosch and Santer 1993). Finally, public health officials have shared anecdotally that adults and copepodites may be more sensitive to Abate™ than nauplii. Critically, the consequences of these stage-structured interactions for disease management are unknown.

To understand copepod demographic responses to episodic culling, and to predict implications for disease control, we conducted a mesocosm experiment that mimicked the 28-day treatment interval of Abate™ by physically removing copepods (either applied equally across stages or biased toward larger individuals). We hypothesized that more intense interventions (higher removal percentages) would cause greater reductions in population density, but that relaxed density dependence might cause compensation or overcompensation between interventions. Further, we speculated that these effects would be more pronounced in populations where both size classes were removed. Finally, we hypothesized that even if copepods rebounded rapidly (‘apparent failure’), risk of parasite transmission might remain low (‘cryptic success’) due to high adult turnover rate and the two-week period required for parasite development from first stage to infectious third stage larvae.

Surprisingly, all interventions failed to reduce mean copepod density over the 14-week intervention period. Copepods recovered extremely rapidly due to relaxed density dependence on maturation rates. However, despite this apparent failure, model simulations indicate that interventions may still reduce transmission risk by specifically suppressing the density of infectious copepods. These results illustrate how density-dependent mechanisms interact with transmission biology to drive cryptic success of control. Stage structure is a key feature of the biology of many pest species, e.g., root-feeding crop pests (Johnson and Rasmann 2015) and mosquito vectors of human diseases (Neale and Juliano 2019, Walker et al. 2021), and control interventions often focus on specific life stages that are not always the proximate cause of harm, e.g., larvicides to control pathogens transmitted by adult mosquitoes (Choi et al. 2019). Thus, understanding the underlying mechanisms of stage-structured density dependence could enhance our ability to address a diverse range of ecological challenges.

## Methods

### Copepod Maintenance and Propagation

We obtained *Eucyclops sp.* copepods isolated near Mandelia, Chad, an active focus of GW transmission. We maintained copepods in 0.5x HHCOMBO artificial lake with 1X micronutrients (hereafter COMBO; Kilham et al. 1998) in 1 L beakers at 26C with a 12:12 daylight cycle. We fed them 7.5 mg/L of powdered egg yolk (Judee’s, Plain City, Ohio) suspension every Monday, Wednesday, and Friday (M/W/F), and transferred copepods to new media approximately every three weeks by sieving. Once densities reached ∼500-1,000 copepods/L, we poured one 1 L culture into 42, 15 L propagation tanks with 4 L COMBO.

### Mesocosm Setup, Sampling, and Experimental Design

On September 20^th^, 2024, we filled 60 15 L tanks with 5 L COMBO and added 1.5 mg/L powdered egg yolk on Monday/Wednesday/Friday to establish microbial communities. On October 2^nd^, 2024 (day 0), we introduced copepods to tanks by adding seven 1 L aliquots collected randomly from the 42 propagation tanks after stirring to homogenize. We then added COMBO to 15 L total and fed tanks 7.5 mg/L every Monday/Wednesday/Friday throughout the experiment. We replaced water weekly to counteract evaporation (<1 L/tank/week).

We maintained the experiment for 126 days and sampled every Friday starting on day 2. To sample, we homogenized tanks (stirring with a plastic paddle) and collected a 500 mL surface grab sample with a beaker. We replaced this volume with COMBO. We intervened at 28-day intervals (days 8, 36, and 64) to mimic the field use of Abate™ (Gindola et al. 2022). We implemented mechanical removal, rather than Abate™, to enable consistent gradients of size-biased and size-unbiased removal. We factorially crossed intervention type (equal vs. size-biased) with the percent of the volume (i.e., copepod population) removed (0-90%: 0, 10, 20, 30, 40, 50, 60, 70, 80, or 90%). For size-biased removal, we sieved water through 100 μm nylon screen, capturing adults, juveniles, and larger nauplii, and returned the water and smaller nauplii to tanks. For size-unbiased (i.e., equal) removal, we replaced the volume removed with fresh COMBO. In Appendix S7, we provide methodological details and results on the accuracy of interventions relative to nominal treatment levels. After the first intervention, we converted the 10% and 20% treatments to 95% and 98% for the following interventions, because low initial intensities caused negligible effects, and higher removal rates better match the stated efficacy of the Abate™ (Grunert et al. 2022). On weeks with interventions, we sampled on Thursday and intervened on Friday. We “skipped” the intervention scheduled for day 92 to observe responses to treatment lapse, as can occur in the field due to inaccessibility from flooding.

### Copepod Counting

We quantified copepods by sieving the 500 mL samples with 35 μm mesh (which captures all stages) and preserving them in 10 mL ethanol. We gently resuspended the samples and examined successive 1 mL aliquots at 10-25x magnification on a dissecting microscope until counting ≥ 50 copepods or screening the entire sample. We counted copepods in three morphologically discernible categories (nauplii, juveniles and eggless adults, and adult females with eggs).

### Statistical Analysis of Intervention Effects on Copepod Density

We conducted all analyses in the R Computing Language v4.3.2 (R Core Team 2023). We tested for differences in mean population density over the first 14 weeks, when treatments were applied “on schedule”, using planned contrasts between all interventions versus controls (Dunnett’s tests; package emmeans (Lenth 2025). This analysis used a Generalized Linear Model (package glmmTMB; Bolker 2017) with mean density as the response, removal proportion, removal type, and their interaction as predictors, and the Gamma error distribution with a log link function.

### State Space Modeling (SSM) of Copepod Population Dynamics

We built a stochastic population dynamic model and simultaneously fit it to all time series of observations from the experiment using the Partially Observed Markov Process (POMP) framework and the panelPomp package (Carles Bretó 2025). We connected likelihood estimates generated by the function pfilter() to an adaptive Markov Chain Monte Carlo (MCMC) sampler in the adaptMCMC package to conduct Bayesian inference, with priors informed by auxiliary experiments (Appendix S2: Figure S2). POMP models are ideal for population dynamics data (Newman et al. 2023) where state variables (the “true” abundances of nauplii, juveniles, and adults) are partially observed as empirical subsample counts of nauplii (N_obs_), juveniles/adult males/adult females with no eggs (JOA_obs_ = “juveniles and other adults”), and adult females (AF_obs_) (Carles Bretó 2025).

We implemented the model by specifying ordinary differential equations that describe deterministic rates of change in the densities of nauplii, *N*, juveniles, *J*, and adults, *A*. In this model (Equations 1-6), nauplii are born, develop into juveniles, die from background causes, and are cannibalized by adults. Juveniles mature from nauplii, develop into adults, and die of background causes. Adults mature from juveniles, birth and cannibalize nauplii, and die of background causes. Critically, birth rate (*b)*, stage-specific death rates (*d_i_,* with *i = N, J, or A*), and stage-specific maturation rates (*m_i_,* with *i = N or J*) depend exponentially on density, with each stage potentially contributing differentially to density dependence via their impact on the competitive environment (i.e., *c_J_* and *c_N_* represent the competitive effects of juveniles and nauplii relative to adults). We allowed adults to cannibalize nauplii following a linear functional response with attack rate α. On intervention days, we incorporated removal as additional mortality on relevant stage(s). During all interventions, we assumed juveniles and adults were removed according to treatment. We assumed nauplii were removed proportionally in size-unbiased removal treatments and used a prior informed by an auxiliary experiment that showed size-biased sieving removed approximately 45% of nauplii (Appendix S2: Methods S1).

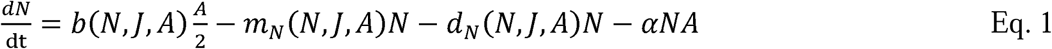

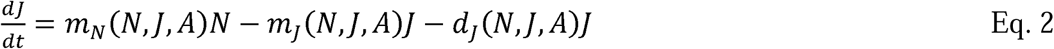

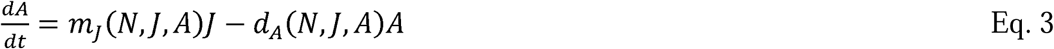

with density-dependent functions:

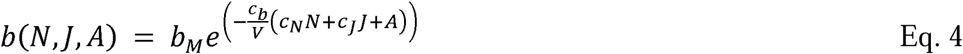

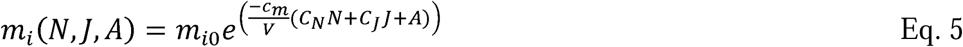

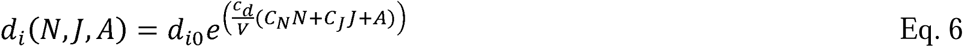

To simulate the dynamics of the latent (or “true and unobserved”) state of the system we assumed demographic stochasticity and implemented the model with a one-day time step. Specifically, we tracked individuals transitioning through stages or dying using random multinomial draws where the probability of three potential outcomes for any individual (remaining in the same state, maturing to the next state, or dying) depends on the relative rates of each process. We assumed births followed Poisson random draws depending on the total birth rate. Our observations differentiate N_obs_, AF_obs_, and JOA_obs_. We assumed that each observation followed a negative binomial distribution with a common overdispersion parameter. For each observation, we assumed the predicted count was determined by the product of the abundance of individuals of that stage at time t, the proportion of the tank sampled (always 0.5/15 L = 3.3%), and the proportion of the 10 mL subsample counted. To align true and observed categories, we assumed a 50:50 sex ratio with female reproduction, with adult females releasing eggs every third day and appearing eggless for one day in each three-day interval (personal observation TDH). With this model specified, we used the pfilter() function in panelPomp to estimate model likelihoods from the data. To maintain variance in log-likelihoods < 1 (Endo et al. 2019), we used 20 particles per likelihood estimation, resulting in variances of approximately 0.5. We ran 32 MCMC chains, with dispersed initial parameter sets for 250,000 iterations each, permitting adaptation during the first 50,000 iterations (burn-in), which we later discarded.

### Model-derived estimates of density dependence

We used model-derived functions of density dependence, observed densities, and stage structure from the experiment to visualize how the competitive environment shaped demographic rates. We thinned chains to every 1,000^th^ iteration, and calculated per-capita birth, death, cannibalism, and maturation rates across a density gradient from 0 to 6,000 individuals/L for each parameter set (the density range observed in the experiment). We fixed stage structure along this density gradient (5.5% adults, 39% juveniles, and 55.5% nauplii, 1:7:10 ratio), based on the average proportions in the experiment. We then generated the mean and 95% CI interval of posterior predictions for each demographic rate along the gradient.

### Compartmental (ODE) stage structured model with interventions and infection

Large-scale experiments with viable GW larvae are currently infeasible. Therefore, we extended our model to gain applied insights into disease control. The extended model (Eq. S1-S11 in Appendix S1: Methods and Equation S1) allows parasites to infect adult copepods. We implemented a distributed developmental delay (Hurtado and Kirosingh 2019) for copepods transitioning from exposed to infectious to definitive hosts (Appendix S1: Methods and Equation S1). This approach uses the “linear chain trick” (Hurtado and Kirosingh 2019) to implement a variable time lag, as we hypothesized this delay could be important for differentiating successful population regulation from successful disease control. Once an adult copepod consumes the parasite, it takes approximately 15 days to become infectious (Lindquist and Cross 2017). Thus, we added a developmental delay of approximately 15 days by implementing a chain of 60 exposed stages that copepods transition through at a rate of 4.3 d^-1^, yielding a 15.45 ± 1.82-day delay from exposure to infectiousness.

Estimates of the force of infection for copepods are unavailable. Therefore, we assumed a 5% infection prevalence among adults, consistent with the limited field data available (Lyons 1972, Steib and Mayer 1988, Garrett et al. 2020), corresponding with λ = 0.0034/day. Further, while parasites are likely introduced in discrete pulses (a single mature female can introduce 1 million L1 larvae (Greenwood 2012)), we implemented a constant force of infection to avoid confounding results with day of introduction. Lastly, we assumed infected adults cannot reproduce (Moorthy 1938).

Using this model, we evaluated the consequences of episodic mortality events on stage-specific and total abundance, with emphasis on the density of infectious copepods, the proximate hazard to definitive hosts. We plotted the temporal dynamics for the 90% mortality scenario, a previously used metric for Abate^TM^ success (South Sudan Guinea Worm Eradication Program 2013). We plotted mean densities of all stages across the whole simulation period for scenarios ranging from 0-90% mortality.

## Results

### Copepod population dynamics and mortality events

Regardless of removal intensity, we found no significant decreases in mean population density over the first 14-weeks of the study for size-unbiased (light blue) or size-biased (dark blue) removal (Fig. 1). This apparent failure was consistent for both the total population and when broken down by stage (Fig. 1, diamond, triangle, circle; all Dunnett’s test *P* > 0.05).

**Figure 1.**
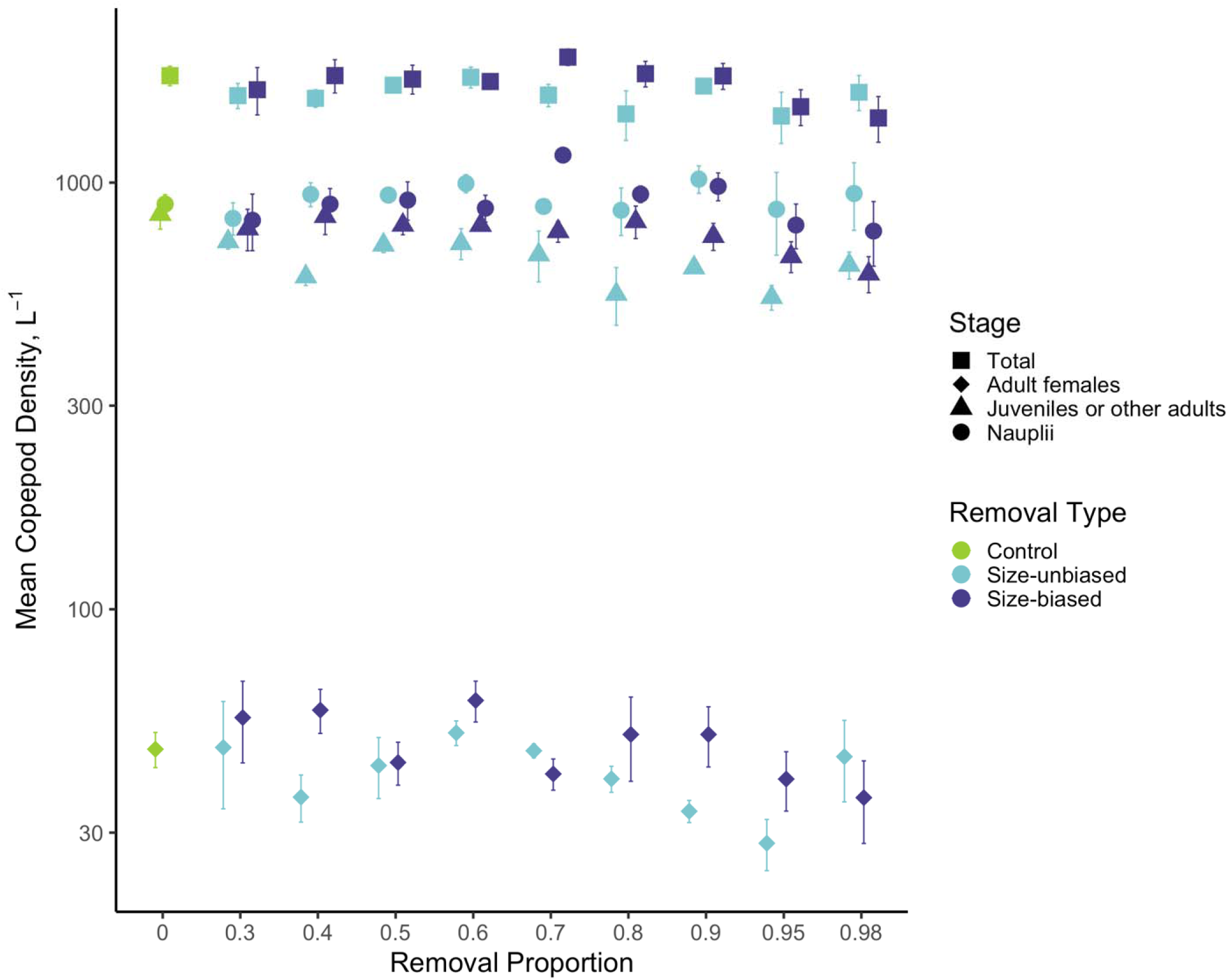
Mean observed copepod densities in the experiment over the 14 weeks in which the interventions were applied on schedule. Points and error bars represent treatment means ± SEs (log-transformed). Regardless of treatment type or removal proportion, there were no significant reductions in total copepod density (squares), or stage-specific densities of egg-bearing adult females (AF; diamonds), juveniles plus other adults (JOA; triangles), or nauplii (N; circles). Similar dynamics were observed regardless of whether removal was selective for large-bodied individuals (size-biased; dark blue) or equally applied across all stages (size-unbiased; light blue).

While we found no effect on mean density, the interventions did reduce densities immediately after application (e.g., 95 and 98% removal, Fig. 2, Appendix S5: Figure S1A-B). However, populations then rapidly rebounded. Populations in controls started at ∼500-1,000 copepods/L, grew slowly for several weeks, and ended with a boom and bust (Fig. 2). In other cases, density in culled tanks temporarily surpassed controls, indicating overcompensation. For example, following the third intervention, densities in the 50% size-biased treatment were briefly higher than controls. In some cases, population recovery occurred so rapidly that declines were imperceptible one week later (e.g., after the second intervention with 80% removal). The best fit of the model generally captured the dynamics in intervention tanks better than that of the control treatment, which is partially expected, as the control treatment represented only 10% of the experiment.

**Figure 2.**
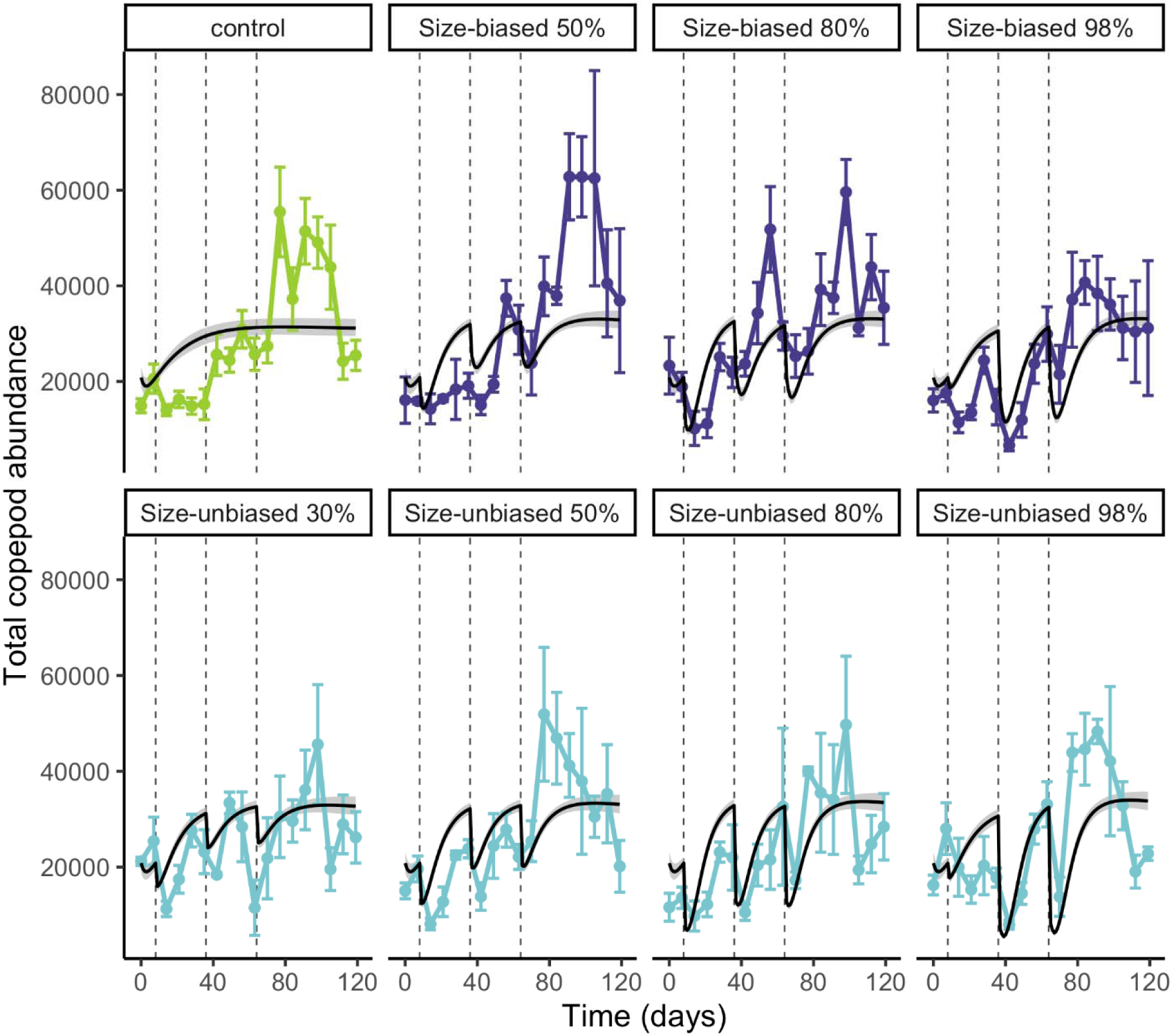
Observed copepod population dynamics (points and error bars represent means ± SE) and best-fitting posterior predictions with 95% credible intervals (black lines and shaded grey regions). Treatments that are representative of the major population dynamics are shown, with all treatments can be found in the Appendix (Appendix S5: Figure S1A). Dynamics in control tanks are plotted in green, size-biased removal tanks are plotted in light blue, and size-unbiased removal tanks are in dark blue. Dashed vertical lines represent interventions. In general, interventions caused immediate reductions in copepod abundance, but populations rebounded rapidly. In some case, e.g., size-biased 50% and size-unbiased 90%, these rebounds surpassed abundances observed in controls. For the 98% treatments, the first intervention was 20% removal, explaining the small dip in the model fit.

### Size-biased Removal

Relative to controls, the 30% and 60% size-biased treatments grew steadily throughout the entire experiment (Fig. 2, Appendix S5: Figure S1A). Populations quickly recovered from interventions, and observed dynamics generally fell within 95% credible intervals of the fitted model predictions. The 40%, 50%, and 70% removal treatments more closely resembled the controls, with large fluctuations in density at the end of the experiment. The 80%, 90%, and 95% removal treatments had multiple extreme booms and busts. Finally, 98% removal treatments fluctuated with low levels of population growth. These shifts sometimes fell outside the credible interval of posterior predictions.

### Size-unbiased Removal

In contrast with controls, 30%, 70% and 80% size-unbiased removal treatments maintained relatively constant abundances across all eighteen weeks, mostly within the credible interval (Fig. 2, Appendix S5: Figure S1A). The 40% and 60% tanks resembled the controls. The 50% treatment also resembled the control; however, instead of final booms and busts, populations declined steadily. The 90%, 95%, and 98% treatments had sustained peaks and troughs, encompassed by the credible interval for most of the experiment. The last few weeks had more extreme booms and busts, not captured within the credible interval.

### Evaluating the Functional Forms of Density Dependence

The strength of negative density dependence varied significantly across processes of birth, maturation, background death, and cannibalism. Density dependence was greatest for maturation rates, with a >18,000-fold reduction across the observed density gradient (Fig. 3). Density dependence on background death rate and cannibalism were also substantial, with a combined 4.4-fold increase over the density range (Fig. 3). However, we estimated essentially no effect on birth rates, with <2% decrease over the density range.

**Figure 3.**
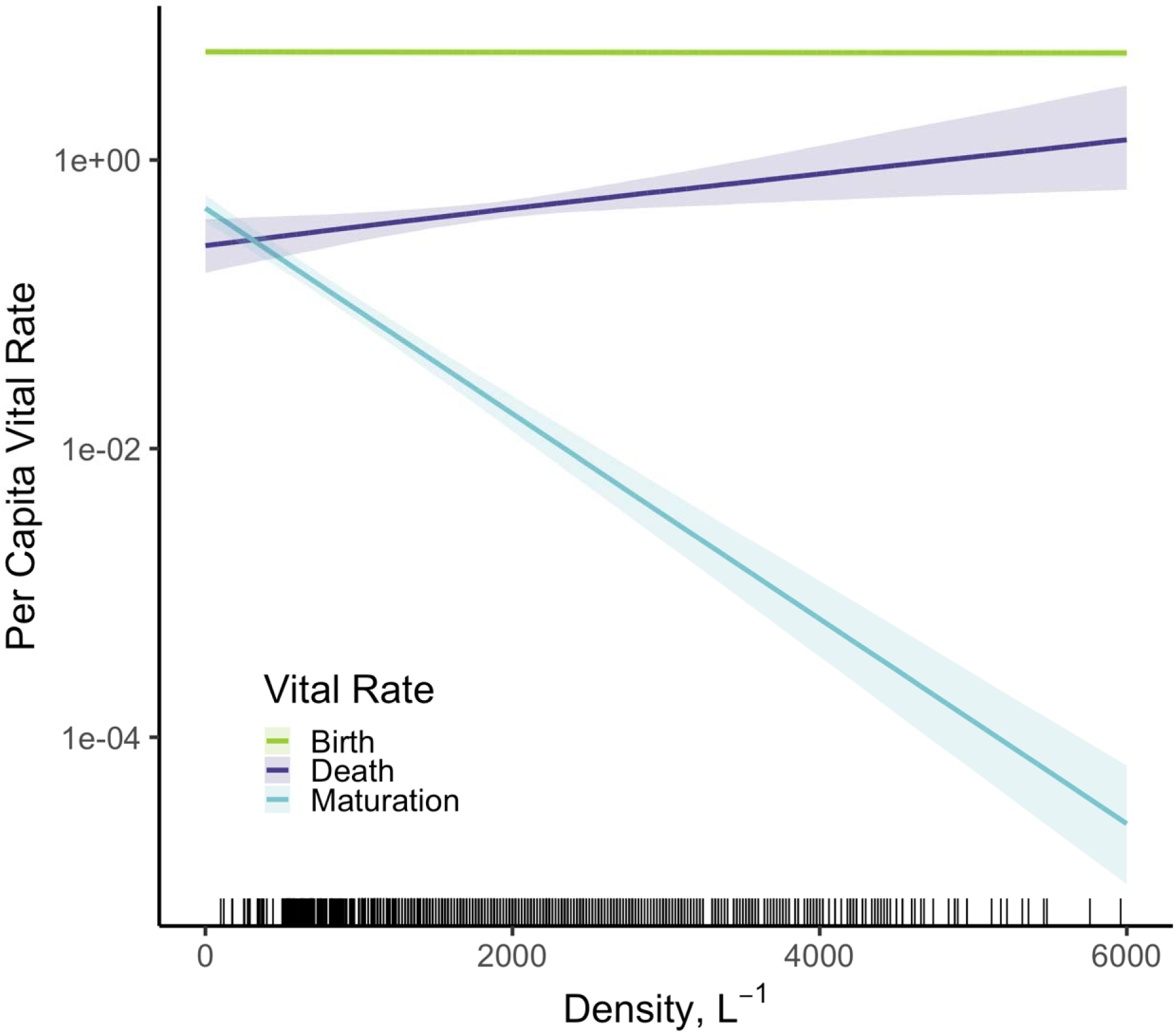
Representation of the sources of negative density dependence affecting copepod populations subject to control interventions. Lines and shaded regions represent mean and 95% credible intervals for posterior model predictions for the effects of copepod population density (with fixed stage structure) on key vital rates: birth rate (green), total death rate of nauplii (dark blue; incorporating background deaths and cannibalism via adults), and naupliar maturation rate (light blue). The rug indicates the observations of population density in the experiment. Adult birth rate was insensitive to competition, declining only 0.9% over the observed density gradient (indicated by the rug along the x-axis). In contrast, death rate increased 4.4-fold over the density gradient. Most importantly, maturation rate was extremely sensitive, declining >18,000 fold over the gradient.

### Transmission Scenario Simulation Results

Simulated GW infection scenarios exhibited rapid, stage-specific recovery of copepods following interventions (Fig. 4). With 90% intervention intensity, nauplii and non-infectious adults rapidly rebounded to densities that surpassed those observed for untreated populations (Fig. 4C, D), and juveniles reached similar, though lower densities until the next intervention (Fig. 4B). Size-unbiased removal generally reduced copepod populations more strongly, but these populations also showed slightly greater overcompensation (Fig. 4A, C). This transient overcompensation reiterated a central result from the experiment (Fig. 1): that mean copepod density appeared unaffected by interventions.

**Figure 4.**
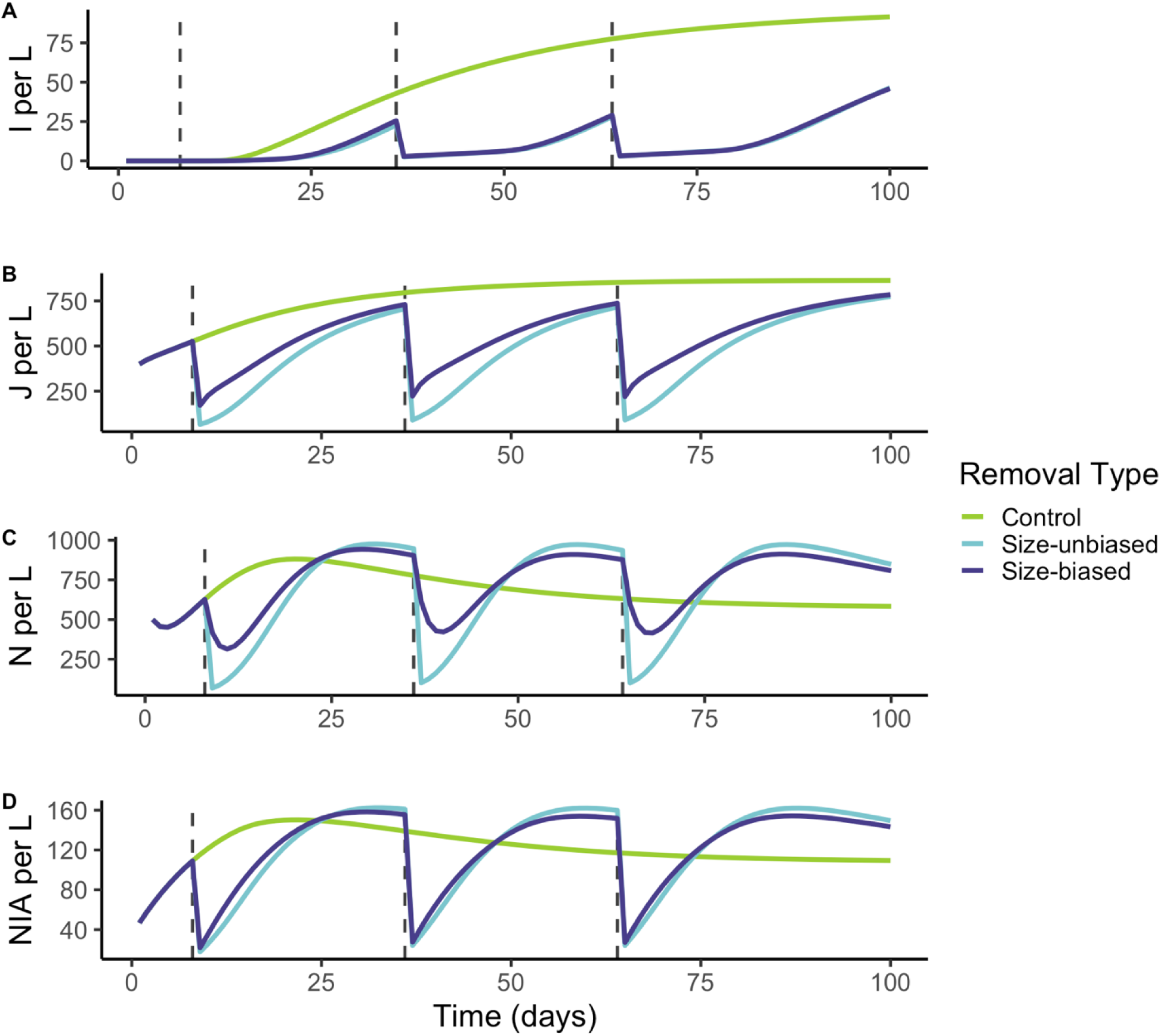
Temporal dynamics from the modeled transmission scenarios incorporating 90% mortality events at 28-day intervals. Dashed vertical lines represent interventions. Copepod stages are shown in different colors. Stages are divided into infectious adults (I), juveniles (J), nauplii (N), and non-infectious adults (NI = susceptible + exposed adults). Control tanks are dashed lines, 90% size-unbiased removal are solid lines, and 90% size-biased removal are the dotted lines. The force of infection parameter (A =0.0034) corresponded with a ∼5% infection rate among adults(Lyons 1972, Steib and Mayer 1988, Garrett et al. 2020). Nauplii and non-infected adults surpass densities of controls (green) with both forms of removal (size-biased [dark blue] and size-unbiased [light blue]). Juveniles nearly recover to densities akin to controls. Most importantly, infectious copepods are severely depressed by both removal types.

Despite this apparent failure, simulated interventions successfully reduced the density of infectious copepods (Fig. 4A). Without intervention, control populations began with low densities of infectious hosts, but these grew steadily (>75 infected copepods L^-1^ by the end of the 100-day simulation). However, both size-unbiased and size-biased suppressed infectious copepod density below 10 L^-1^ (Fig. 4A). Evaluating mean densities of stages across an intervention intensity gradient (Fig. 5), total population density and non-infected adults (susceptible + exposed classes) remained relatively constant at all intensities, while the density of infectious adults steadily declined across the gradient. Thus, despite observing that copepod populations rebound rapidly – enabled by intense density dependence on maturation rates – the density of infectious hosts, the proximate source of mammalian infections, is highly sensitive to intervention.

**Figure 5:**
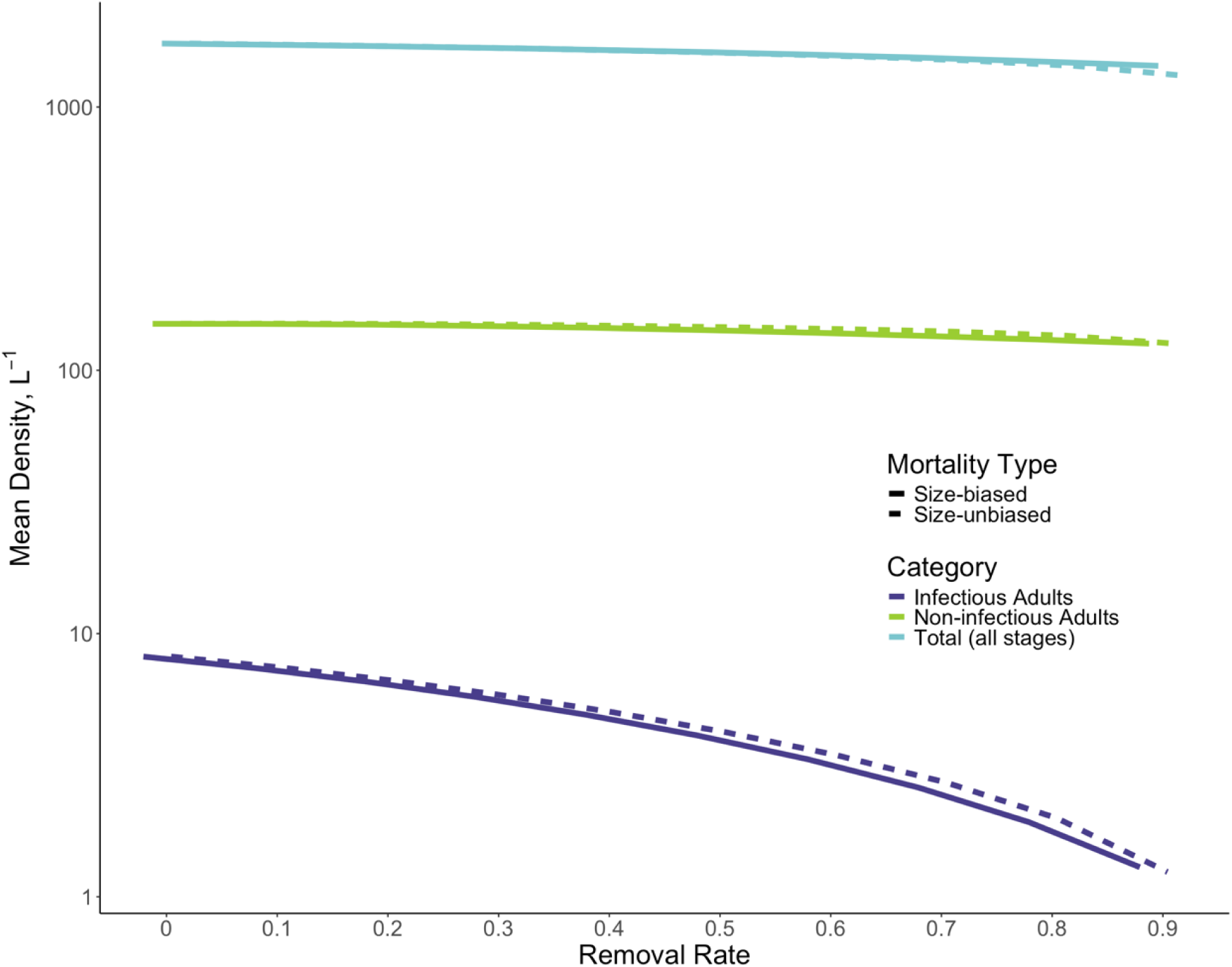
Mean total density of copepod populations (light blue), non-infectious adults (green), and infectious adults (dark blue) in 120-day transmission scenarios modeled along a gradient of intervention intensity for size-biased (solid line) and size-unbiased (dashed line) strategies. As in the experiment, total density and the density of non-infectious adults (susceptible and exposed) were largely insensitive to interventions, remaining relatively constant despite 90% population reductions imposed every 28 days. In contrast, the mean density of infectious adults, the proximate driver of mammalian hazard of infection, decreased almost 10-fold along the same gradient. Thus, despite the rapid recovery of copepod populations and the appearance of failed control, intervention can be highly effective at reducing the proximate driver of infection in humans and other mammals.

## Discussion

Uncovering the mechanisms regulating populations is a fundamental goal in population ecology (Kaspersson et al. 2013, Karatayev et al. 2015). We combined a pest control experiment and mechanistic modeling to partition the magnitude and importance of four general putative mechanisms of density regulation. Then, we integrated these empirically-derived regulating effects with a transmission model to evaluate how relaxation of density dependence via realistic control methods shapes effectiveness of interventions for GW, a target for global parasite eradication (Pellegrino et al. 2022).

Our experiment investigated how copepod populations – intermediate hosts of GWD – responded to periodic culling. We designed treatments to mimic the treatment of water sources with Abate™, an intervention commonly used to prevent GWD in endemic countries. We hypothesized that more aggressive control efforts would better suppress copepod populations, but that relaxed density dependence might lead to overcompensation. Strikingly, even when we episodically removed up to 98% of the population, densities across were statistically indistinguishable from controls (Fig. 1, Appendix S3: Table S1). Similar rebounds have been observed in other copepod species, including *Mesocyclops leuckarti* which overshot pre-treatment densities following application of Triphenyltin, a fungicide (Kulkarni et al. 2014). Our interventions had failed to reduce copepod densities, an alarming prospect as other apparent pest control failures have posed immense challenges for public health and conservation practitioners (Rivero et al. 2010, Neale and Juliano 2019, Grosholz et al. 2021). For example, populations of mosquitoes, sand flies, assassin bugs, and fleas, which are routinely treated to control diseases such as malaria, leishmania, chagas, and plague, respectively, show evidence of insecticide resistance, threatening vector control (Cao et al. 2025). Further, specific mosquito species including *Aedes* and *Culex sp.*, vectors of diseases such as dengue, chikungunya, Zika virus (*Aedes*) (Patterson et al. 2016), and West Nile Virus (*Culex*) (Hamer et al. 2008), show evidence of culling-induced overcompensation, indicating that control strategies must be careful to not impose mortality at levels that lead to overcompensatory releases from density dependence (Neale and Juliano 2019). A striking example of a pest control program that backfired is the effort to eliminate the invasive European green crab in an estuary off of California (Grosholz et al. 2021). Culling of green crabs was an effective strategy for years until the population density was reduced below a critical threshold, triggering stage-specific overcompensation in recruitment (Grosholz et al. 2021). The result was a boom in juvenile crabs, a shift in size structure, and exacerbation of negative impacts (Grosholz et al. 2021). Together, these examples underscore the challenges of pest control – especially in light of stage structure.

To attempt to further explain the surprisingly fast recovery we observed, we investigated the forms of density dependence regulating *Eucyclops sp.* populations. Few studies have parsed specific drivers of density dependence in natural populations. In one example, stocking large brown trout reduced the growth rate of smaller individuals (Kaspersson et al. 2013). In another, dynamics of Swedish vole populations were consistent with density dependence in recruitment alone (Lambin et al. 2025). Stage-specific density dependence in maturation rates has been documented in some ecological systems (Karatayev et al. 2015, Walker et al. 2021), including other copepods (Kulkarni et al. 2014), allowing for population persistence under significant mortality (Karatayev et al. 2015). We determined that density dependence in *Eucyclops sp.* was driven predominantly by maturation rates, with a >18,000-fold reduction along the density gradient, with a modest ∼4.4-fold increase in total mortality, and no change in birth rate (Fig. 3). Thus, when interventions occur, surviving adults maintain birth rates, but nauplii mature extremely quickly. Consequently, higher proportions of nauplii and juveniles survive to development, enabling rapid recovery and temporary overcompensation. (Figs. 2 and 4).

For the goal of GWD management, the failure to reduce mean copepod population densities seems alarming. However, not all copepod life stages host GW, (McCullough 1983, Bimi 2007) and there is a significant delay between copepod exposure to GW and infectiousness to mammals (∼15 days; Gonzalez Engelhard et al. 2021). Thus, we speculated that if population rebounds were driven by overcompensation in young stages that cannot transmit GW, then apparent failures to control the overall population might actually conceal successes. We evaluated this possibility by extending our model to include GW infection of adults with a distributed delay from the exposed to infectious class. We tracked changes in the density and stage structure of copepod populations across a gradient of intervention intensity.

When we simulated GW transmission scenarios, we found strong support for stage-specific recovery dynamics, as documented in other zooplankton systems (Nilsson et al. 2010, Kulkarni et al. 2014). Specifically, the densities of nauplii, juveniles, and non-infectious adults all recovered rapidly to meet or exceed pre-intervention densities following each successive mortality event (Fig. 4), indicating compensation or overcompensation (Abrams 2009, Kulkarni et al. 2014, de Roos 2020, Grosholz et al. 2021). Critical to GW transmission, the density of infectious adults remained extremely low. Thus, despite the apparent failure of these interventions to control copepod populations, they appear highly effective at specifically controlling the life stage that is the proximate cause of GW infections. We attribute this effect to the approximately15-day developmental period from first stage to infectious third stage GW larvae (Lindquist and Cross 2017, Gonzalez Engelhard et al. 2021), which is long enough to prevent accumulation of infectious copepods before each successive intervention (Fig 4., Fig. 5). Similar lags of parasite development in intermediate hosts and vectors are present in other systems. For example, in Anopheline mosquitoes, a 2-3 week incubation period separates vector exposure to *Plasmodium* and infectiousness to humans (Crutcher JM 1996). Similarly, *Trypanosoma* maturation in tsetse flies requires approximately three weeks (Telleria et al. 2014). Thus, reducing the overall density of intermediate hosts or vectors may not be essential for success if control can cause high turnover in the stage(s) responsible for transmission.

Our results raise the importance of field surveillance targeted at infected copepods. At present, diagnosing infections in copepods from field is infeasible due to GW rarity and the scale of geographic surveillance (Pellegrino et al. 2022). Therefore, current eradication programs rely on monitoring the total density of copepods before and after Abate^TM^ applications. If copepod communities in the field recover as rapidly as the *Eucyclops sp.* examined in our study, officials might falsely conclude that control is ineffective, because population densities return to pre-treatment levels every month. However, our experiment and model suggest that this observation could conceal a cryptic success: reduction in the density of infectious copepods. Thus, we recommend the development of tools to facilitate large-scale spatial and temporal diagnostics of GW-infected copepods to evaluate the relevance of this counterintuitive finding.

We note some important caveats to translating these insights to the field. First, we focused on a single copepod isolate. Greater understanding of copepod diversity in demographic rates and sensitivity to Abate^TM^ would greatly benefit eradication programs. Second, we implemented mechanical removal, which enabled a precise experimental design, but can differ from chemical control. Experiments with Abate^TM^ are needed to evaluate chemical persistence, delayed or sublethal effects, and the consequences of over- or underdosing. Lastly, our analysis identified high importance for negative density dependence on maturation rates and mortality, and no effect on birth rates. Generally, the model fit the data well for the first 70-80 days of the experiment, albeit with some initial lags in population growth (Fig. 2). This imperfect fit could be due to missing key processes in the model, such as an Allee effect, where mating failure occurs at low population densities (seen for copepods (Kramer et al. 2008)). The inability of the model to precisely recover empirical dynamics could also be due to nonconstant conditions in the tanks (e.g., the accumulation of nutrients (Sahandi et al. 2023) and/or waste products or differences in temperature or photoperiod over the course of the experiment). Nevertheless, the POMP approach has major benefits, including representation of constraints in the observation process and incorporation of process noise. Lastly, future studies of copepod control and recovery could consider other functional forms for key demographic processes, including Type-II or Type-III functional responses for cannibalism.

Chemical control of copepod populations represents a critical intervention for national GW eradication programs (Gindola et al. 2022). This study highlights that nuance may be needed to evaluate this intervention: population dynamics may suggest that control efforts are failing or even backfiring, but direct analyses of infectious copepods, the proximate cause of human cases and animal infections should be prioritized. Future laboratory studies should expand to greater copepod diversity and directly evaluate pesticide formulations in use. Ultimately, a deeper understanding of copepod control may improve the portfolio of GWD interventions and contribute to the eradication of this devastating parasite.

More broadly, pest control is inherently designed to reduce population density (Ni et al. 2025), so understanding density dependence is critical to predicting responses (McIntire and Juliano 2018). This work underscores the value of defining which life stages of pests are most impactful, elucidating how stage-structured pest populations respond to interventions, and developing a mechanistic understanding of density dependent mechanisms regulating populations under control, especially in cases where focal populations experience drastic and rapid shifts in density. Together, this information can be applied to anticipate and explain potentially counterintuitive consequences of interventions. A thorough understanding of these dynamics is critical for forecasting long-term population trajectories and their sensitivity to density-dependent perturbations, paramount to predicting the influence of environmental and ecological stressors on population dynamics.

## Acknowledgements

This work was supported by funding from The Carter Center. We thank the Programme National d’Eradication du Ver de Guinée - Tchad for collecting and providing the Chadian *Eucyclops sp.* copepods used in our study, and the Yen lab at the Georgia Institute of Technology for providing the culture to us. We thank The Carter Center and other Civitello lab members for feedback on this manuscript. We also thank M. Hoogshagen and J. Raytselis for helping with sampling over the holidays, and B. Lukubye for moral support.

## Author Contributions

N.R., S.G., A.S., and D.J.C. developed and planned experiments. N.R., M.R., L.S., L.T.S, and J.M. conducted the experiment. N.R., M.R., L.S., L.T.S, J.M., T.H., E.R., and I.G., processed samples. N.R., K.X., and D.J.C. conducted analysis. A.S. and D.J.C. provided feedback to refine analysis. N.R. wrote manuscript and made figures with feedback from A.S. and D.J.C..

## Literature Cited

Abrams, P. A. 2009. When does greater mortality increase population size? The long history and diverse mechanisms underlying the hydra effect. Ecology Letters 12:462–474.

Bimi, L. 2007. Potential vector species of Guinea worm (Dracunculus medinensis) in Northern Ghana. Vector Borne Zoonotic Dis 7:324–329.

Bolker, B. 2017. {glmmTMB} Balances Speed and Flexibility Among Packages for Zero-inflated Generalized Linear Mixed Modeling.

Calizza, E., M. L. Costantini, G. Careddu, and L. Rossi. 2017. Effect of habitat degradation on competition, carrying capacity, and species assemblage stability. Ecology and Evolution 7:5784–5796.

Cao, L. J., J. C. Chen, J. A. Thia, T. L. Schmidt, R. Ffrench-Constant, L. X. Zhang, Y. Yang, M. C. Yuan, J. Y. Zhang, X. Y. Zhang, Q. Yang, Y. J. Gong, H. Li, X. Chen, A. A. Hoffmann, and S. J. Wei. 2025. Recurrent mutations drive the rapid evolution of pesticide resistance in the two-spotted spider mite Tetranychus urticae. Elife 14.

Carles Bretó, J. W., Aaron A. King, and Edward L. Ionides. 2025. panelPomp: Analysis of Panel Data via Partially Observed Markov Processes in R. The R Journal.

Choi, L., S. Majambere, and A. L. Wilson. 2019. Larviciding to prevent malaria transmission. Cochrane Database Syst Rev 8:Cd012736.

Crutcher JM, H. S. 1996. Malaria, Medical Microbiology.

de Roos, A. 2020. The impact of population structure on population and community dynamics. Pages 53–73.

Dennis, B., R. A. Desharnais, J. M. Cushing, S. M. Henson, and R. F. Costantino. 2001. Estimating Chaos and Complex Dynamics in Insect Populations Ecological Monographs 71:277–303.

Endo, A., E. van Leeuwen, and M. Baguelin. 2019. Introduction to particle Markov-chain Monte Carlo for disease dynamics modellers. Epidemics 29:100363.

Garrett, K. B., E. K. Box, C. A. Cleveland, A. A. Majewska, and M. J. Yabsley. 2020. Dogs and the classic route of Guinea Worm transmission: an evaluation of copepod ingestion. Sci Rep 10:1430.

Gindola, Y., D. Getahun, K. Mohammed, E. Kamau, B. Camara, M. Wossen, K. Demissie, S. Abdela, G. Gebrewolde, G. Hailu, M. Tegistu, A. Okugn, and G. Gikilo. 2022. Abate application practices in the Guinea worm endemic region of Gambella, Ethiopia: identification of elimination gaps. The Journal of Infection in Developing Countries 16:20S–25S.

Gonzalez Engelhard, C. A., A. P. Hodgkins, E. E. Pearl, P. K. Spears, J. Rychtář, and D. Taylor. 2021. A mathematical model of Guinea worm disease in Chad with fish as intermediate transport hosts. Journal of Theoretical Biology 521:110683.

Gotelli, N. 2008. A Primer of Ecology Fourth edition.

Greenwood, D. 2012. 63 - Helminths: Intestinal worm infections; filariasis; schistosomiasis; hydatid disease. Pages 655–666 *in* D. Greenwood, M. Barer, R. Slack, and W. Irving, editors. Medical Microbiology (Eighteenth Edition). Churchill Livingstone, Edinburgh.

Grosholz, E., G. Ashton, M. Bradley, C. Brown, L. Ceballos-Osuna, A. Chang, C. de Rivera, J. Gonzalez, M. Heineke, M. Marraffini, L. McCann, E. Pollard, I. Pritchard, G. Ruiz, B. Turner, and C. Tepolt. 2021. Stage-specific overcompensation, the hydra effect, and the failure to eradicate an invasive predator. Proceedings of the National Academy of Sciences 118:e2003955118.

Grunert, R., E. Box, K. Garrett, M. Yabsley, and C. Cleveland. 2022. Effects of Temephos (Abate®), Spinosad (Natular®), and Diflubenzuron on the Survival of Cyclopoid Copepods. Am J Trop Med Hyg 106:818–822.

Hamer, G. L., U. D. Kitron, J. D. Brawn, S. R. Loss, M. O. Ruiz, T. L. Goldberg, and E. D. Walker. 2008. Culex pipiens (Diptera: Culicidae): A Bridge Vector of West Nile Virus to Humans. Journal of Medical Entomology 45:125–128.

Hanazato, T., T. Iwakuma, M. Yasuno, and M. Sakamoto. 1989. Effects of temephos on zooplankton communities in enclosures in a shallow eutrophic lake. Environmental Pollution 59:305–314.

Hixon, M. A., and G. P. Jones. 2005. Competition, Predation, and Density-Dependent Mortality in Demersal Marine Fishes Ecology 86:2847–2859.

Hurtado, P. J., and A. S. Kirosingh. 2019. Generalizations of the ‘Linear Chain Trick’: incorporating more flexible dwell time distributions into mean field ODE models. J Math Biol 79:1831–1883.

Johnson, S. N., and S. Rasmann. 2015. Root-feeding insects and their interactions with organisms in the rhizosphere. Annu Rev Entomol 60:517–535.

Karatayev, V. A., C. E. Kraft, and E. F. Zipkin. 2015. Racing through life: maturation rate plasticity regulates overcompensation and increases persistence. Ecosphere 6:art203.

Kaspersson, R., F. Sundström, T. Bohlin, and J. I. Johnsson. 2013. Modes of Competition: Adding and Removing Brown Trout in the Wild to Understand the Mechanisms of Density-Dependence. PLOS ONE 8:e62517.

Kilham, S., D. Kreeger, S. Lynn, C. Goulden, and L. Herrera. 1998. COMBO: A defined freshwater culture medium for algae and zooplankton. Hydrobiologia 377:147–159.

Kramer, A. M., O. Sarnelle, and R. A. Knapp. 2008. Allee Effect Limits Colonization Success of Sexually Reproducting Zooplankton. Ecology 89:2760–2769.

Kulkarni, D., U. Hommen, A. Schäffer, and T. G. Preuss. 2014. Ecological interactions affecting population-level responses to chemical stress in Mesocyclops leuckarti. Chemosphere 112:340–347.

Lambin, X., M. Begon, S. J. Burthe, I. M. Graham, J. L. MacKinnon, S. Telfer, and M. K. Oli. 2025. Density-dependent recruitment but not survival drives cyclic dynamics in a field vole population. Proceedings of the National Academy of Sciences 122:e2509516122.

Leboulanger, C., C. Schwartz, P. Somville, A. O. Diallo, and M. Pagano. 2011. Sensitivity of Two Mesocyclops (Crustacea, Copepoda, Cyclopidae), from Tropical and Temperate Origins, to the Herbicides, Diuron and Paraquat, and the Insecticides, Temephos and Fenitrothion. Bulletin of Environmental Contamination and Toxicology 87:487–493.

Lenth, R. V. 2025.emmeans: Estimated Marginal Means, aka Least-Squares Means.

Lindquist, H. D. A., and J. H. Cross. 2017. 195 - Helminths. Pages 1763–1779.e1761 *in* J. Cohen, W. G. Powderly, and S. M. Opal, editors. Infectious Diseases (Fourth Edition). Elsevier.

Lu, A., and L. Wang. 2025. Hygiene conditions explain larval density-dependent performance in Plutella xylostella with sufficient food. Journal of Asia-Pacific Entomology 28:102444.

Lyons, G. R. 1972. Guineaworm infection in the Wa district of north-western Ghana. Bull World Health Organ 47:601–610.

McCullough, F. S. 1983. Cyclopoid copepods: their role in the transmission and control of dracunculiasis. World Health Organization, Geneva, Ecology and Control of Vectors Unit, Division of Vector Biology and Control.

McIntire, K. M., and S. A. Juliano. 2018. How can mortality increase population size? A test of two mechanistic hypotheses. Ecology 99:1660–1670.

Moorthy, V. N. 1938. Observations on the development of Dracunculus medinensis larvae in Cyclops. American Journal of Hygiene.

Neale, Z. R., and S. A. Juliano. 2019. Finding the sweet spot: What levels of larval mortality lead to compensation or overcompensation in adult production? Ecosphere 10:e02855.

Newman, K., R. King, V. Elvira, P. de Valpine, Rachel S. McCrea, and B. J. T. Morgan. 2023. State-space models for ecological time-series data: Practical model-fitting. Methods in Ecology and Evolution 14:26–42.

Ni, J., J. Wang, W. Zhang, E. Chen, Y. Gao, J. Sun, W. Huang, J. Xia, W. Zeng, J. Guo, and Z. Gong. 2025. Sustainable control and integrated management through a one health approach to mitigate vector-borne disease. One Health 20:101018.

Nilsson, K. A., L. Persson, and T. Van Kooten. 2010. Complete compensation in Daphnia fecundity and stage-specific biomass in response to size-independent mortality. Journal of Animal Ecology 79:871–878.

Park, T., D. B. Mertz, W. Grodzinski, and T. Prus. 1965. Cannibalistic Predation in Populations of Flour Beetles. Physiological Zoology 38:289–321.

Patterson, J., M. Sammon, and M. Garg. 2016. Dengue, Zika and Chikungunya: Emerging Arboviruses in the New World. West J Emerg Med 17:671–679.

Pellegrino, C., G. Patti, M. Camporeale, A. Belati, R. Novara, R. Papagni, L. Frallonardo, L. Diella, G. Guido, E. De Vita, V. Totaro, F. V. Segala, N. Veronese, S. Cotugno, D. F. Bavaro, G. Putoto, N. Bevilacqua, C. Castellani, E. Nicastri, A. Saracino, and F. Di Gennaro. 2022. Guinea Worm Disease: A Neglected Diseases on the Verge of Eradication. Trop Med Infect Dis 7.

R Core Team. 2023.R: A Language and Environment for Statistical Computing.

Rivero, A., J. Vézilier, M. Weill, A. F. Read, and S. Gandon. 2010. Insecticide Control of Vector-Borne Diseases: When Is Insecticide Resistance a Problem? PLOS Pathogens 6:e1001000.

Sahandi, J., P. Sorgeloos, H. Xiao, F. Mu, X. Wang, Z. Qi, Y. Zheng, and X. Tang. 2023. The use of microbes in culture of harpacticoid copepod (Tigriopus japonicus Mori) for suppression of Vibrio and enhancement of population growth, enzyme activity and microbial longevity. Aquaculture 563:739008.

Sebens, K. P. 1982. Competition for Space: Growth Rate, Reproductive Output, and Escape in Size. The American Naturalist 120:189–197.

Simonetti, O., V. Zerbato, C. Maurel, L. Cosimi, S. Babich, F. Cavalli, S. Di Bella, D. Pavia, C. Pesaresi, and R. Luzzati. 2023. The current state of knowledge on dracunculiasis: a narrative review of a rare neglected disease. Infez Med 31:500–508.

Smith, G. C. 1997. An analysis of the form of density dependence in a simulation model of a seasonal breeder undergoing control. Ecological Modelling 95:181–189.

South Sudan Guinea Worm Eradication Program. 2013. Management and Technical Guidelines.*in* M. o. H.-R. o. S. Sudan, editor.

Steib, K., and P. Mayer. 1988. Epidemiology and vectors of Dracunculus medinensis in northwest Burkina Faso, West Africa. Annals of Tropical Medicine & Parasitology 82:189–199.

Stockseth, L., Z. Neale, and V. H. W. Rudolf. 2025. Strengthening of negative density dependence mediates population decline at high temperatures. Ecology 106:e70030.

Svanbäck, R., and D. I. Bolnick. 2007. Intraspecific competition drives increased resource use diversity within a natural population. Proceedings of the Royal Society B: Biological Sciences 274:839–844.

Telleria, E. L., J. B. Benoit, X. Zhao, A. F. Savage, S. Regmi, T. L. A. e Silva, M. O’Neill, and S. Aksoy. 2014. Insights into the Trypanosome-Host Interactions Revealed through Transcriptomic Analysis of Parasitized Tsetse Fly Salivary Glands. PLOS Neglected Tropical Diseases 8:e2649.

van den Bosch, F., and W. Gabriel. 1994. A model of growth and development in copepods. Limnology and Oceanography 39:1528–1542.

van den Bosch, F., and B. Santer. 1993. Cannibalism in Cyclops abyssorum. Oikos 67:19–28.

Walker, M., M. A. Robert, and L. M. Childs. 2021. The importance of density dependence in juvenile mosquito development and survival: A model-based investigation. Ecological Modelling 440:109357.

World Health Organization. 1986. Decisions and list of resolutions. World Health Assembly.

World Health Organization. 2025. Dracunculiasis (Guinea-worm disease).

