## Appendix for "Apparent failure and cryptic success of disease control via intermediate host population reduction"

In this appendix, we present supplementary materials regarding:

(A) Appendix S1: Ordinary differential equation model with infections and cannibalism (Methods and Equation S1, Figure S2A, Figure S2B)
(B) Appendix S2: Bayesian Priors (Methods S1, Figure S2)
(C) Appendix S3: Parameter estimates (Table S1)
(D) Appendix S4: Posterior distributions (Figure S1)
(E) Appendix S5: Model fit to data (Methods S1, Figure S2A, Figure S2B)
(F) Appendix S6: Testing for density dependence in reproduction (Methods S1, Equation S2, Results S3, Figure S4)
(G) Appendix S7: Evaluating bias and precision of removal copepods removed during interventions. interventions (Methods S1)

**(A) Appendix S1: Ordinary differential equation model with infections and cannibalism**

**Supplementary Methods and Equations S1:** We modified Eq. 1-6 from the main text to include parasite exposure and infection.

$\frac{dN}{\mathrm{dt}}=b\left( N,J,A \right)\frac{A}{2}-m_{N}\left( N,J,A \right)N-d_{N}\left( N,J,A \right)N-\alpha NA$	Eq. S1

$\frac{dJ}{dt}=m_{N}\left( N,J,A \right)N-m_{J}\left( N,J,A \right)J-d_{J}\left( N,J,A \right)J$	Eq. S2

$\frac{dA}{dt}=m_{J}\left( N,J,A \right)J-d_{A}\left( N,J,A \right)A$	Eq. S3

$\frac{dE_{1}}{dt}=\lambda A-l_{R}E_{1}-d_{A}cE_{1}$	Eq. S4

$\frac{dE_{s}}{dt}= l_{R}E_{s-1}-l_{R}E_{s}-d_{A}cE_{s}$	Eq. S5

$\frac{dI}{dt}= {l_{R}E}_{sFinal}-d_{A}cI$	Eq. S6

$b\left( N,J,A \right) = \frac{b_{M}\left( A+sum\left( E_{s} \right) \right)}{2}{\times e}^{\left( -\frac{{comp}_{b}}{V}\left( c_{N}N+c_{J}J+A \right) \right)}$	Eq. S7

$m\left( N,J,A \right)=m \times e^{\left( \frac{-comp_{m}}{VOL}\times(C_{N}\times N+C_{J}\times J+A+sum\left( E_{s} \right)+I) \right)}$	Eq. S8

$m_{N}c$ *=* $m_{J}(N,J,A)$	Eq. S9

$m_{J}c$ *=* $m_{N}(N,J,A)$	Eq. S10

$d\left( N,J,A \right)=d\times e^{\left( \frac{comp_{d}}{VOL}\times(C_{N}\times N+C_{J}\times J+A+sum\left( E_{s} \right)+I) \right)}$	Eq. S11

Specifically, we extended the model by assuming that adult individuals, *A*, become exposed with a constant force of infection, λ. Exposure causes individual adults to transition to the first exposed class, *E_1_*. Thereafter, exposed individuals transition through an arbitrary number of exposed, *E_S_*, classes at a constant rate, *l*, and then finally into the infectious class, *I*. This “linear chain trick”(Hurtado and Kirosingh 2019) results in a distributed delay from exposure to infectiousness that mimics observed data for copepods and GW, latent period = 15.45 ± 1.82 days (mean ± SE, REF).

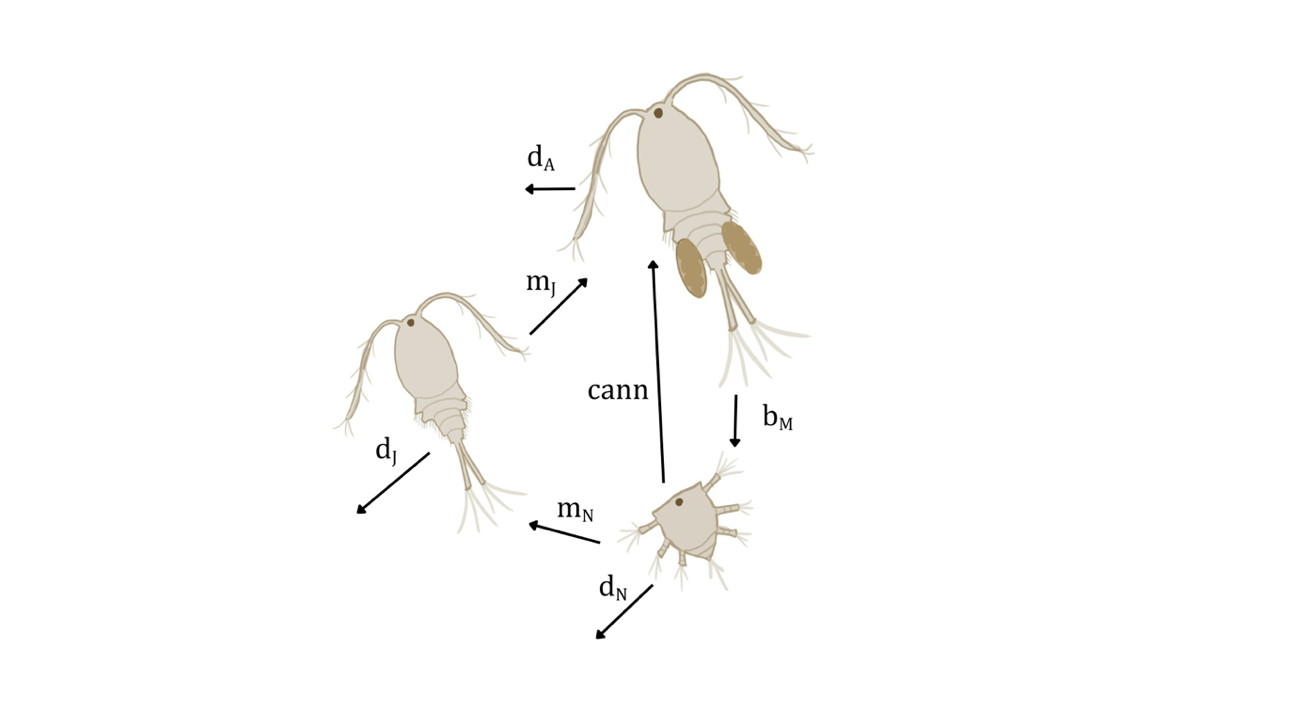

**Figure S2A:** Diagram of key processes in the model fit to the experimental data and represented in Equations 1-6 of the main text. Nauplii are born, cannibalized, die, and mature. Juveniles die or mature. Adults give birth, eat nauplii, and die.

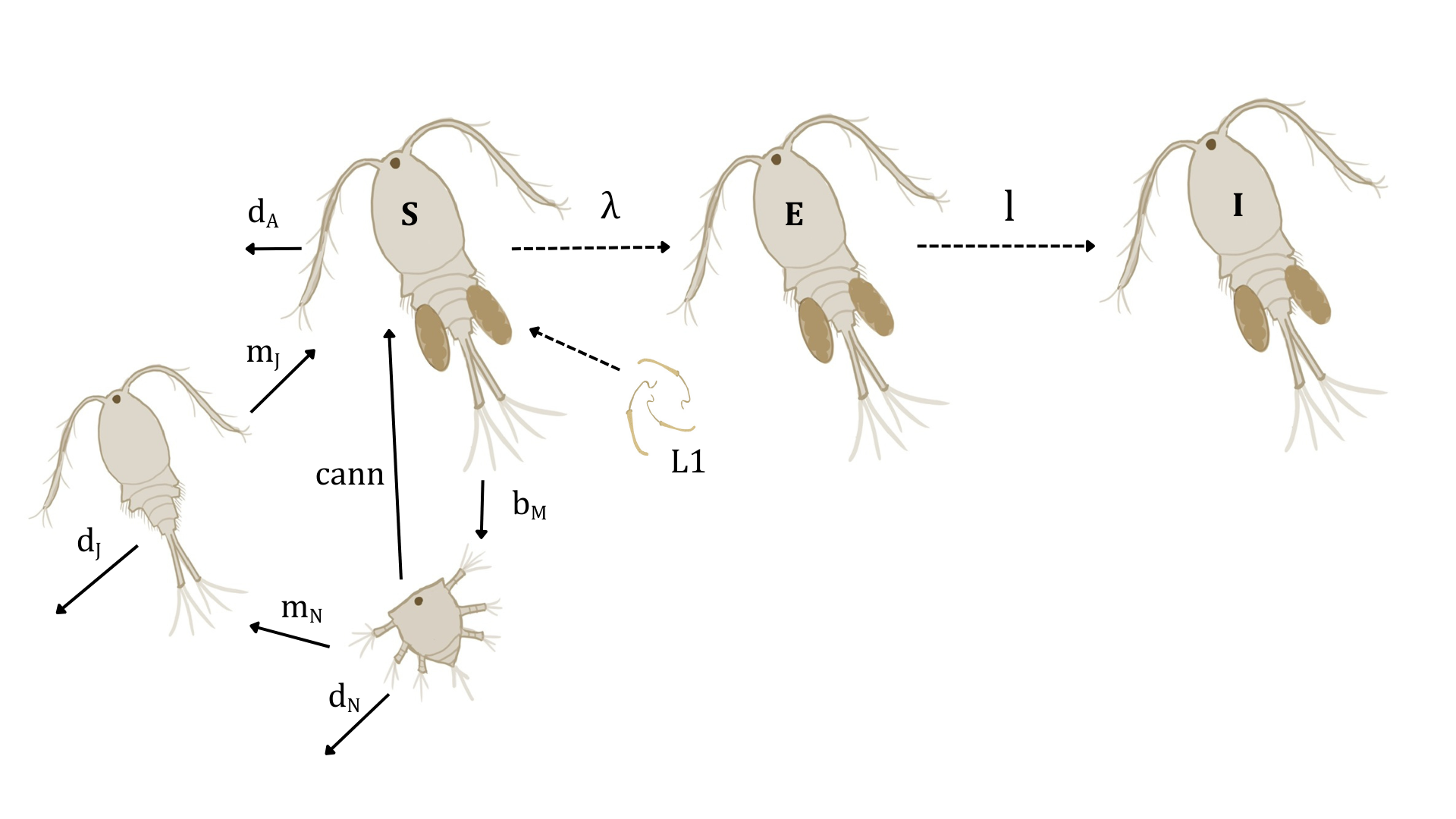

**Figure S2B:** Diagram of the model extension used to simulate GW transmission scenarios and presented in Equations S1-S11 in this supplement. Nauplii are born, cannibalized, die, and mature. Juveniles die or mature. Adults give birth, eat nauplii, die, or become infected. Exposed copepods remain transition through 60 exposed stages with a transition rate, *l* = 4.3 d-1, resulting in a ~15 day latent period.

**(B) Appendix S2: Bayesian priors and estimation of the posterior**

**Methods S1:** We incorporated prior information to constrain some of the parameters that we independently estimated in auxiliary short-term laboratory experiments. We estimated the maturation rate of individuals held at low densities (N = 32 replicate clutches, daily observations over 12 days), yielding estimates of m_N0_ = 0.457 ± 0.11 and m_J0_ = 0.067 ± 0.035 (mean ± SE, units: d^-1^). We estimated maximal birth rate for isolated adult females (N = 12), yielding an estimate of b_M_ = 5.67 ± 0.19 (mean ± SE, units: eggs female^-1^ d^-1^). We implemented these as normally distributed priors. We also incorporated our independent estimate of nauplii removal efficiency, mean = 45.3%, 95% CI 41.9 – 48.7%, as a beta distributed prior with parameters α = 416.25 and β = 508.75. All other parameters received vague uniform priors (if unconstrained) or vague beta priors (α = 1, β = 1) if constrained to the range 0-1. For any given parameter set, we estimated the posterior log-likelihood as the sum of the log-likelihood estimated by the pfilter() function plus the log-likelihood of the prior. We then used the MCMC() function in the adaptMCMC package to conduct implement MCMC via an adaptive Metropolis sampler. We ran 32 chains in parallel on the Emory University Biology Department cluster. Each chain had a 50,000 iteration adaptation phase followed by another 200,000 iteration phase used for parameter estimation. All 32 chains converged to the highest posterior likelihood region.

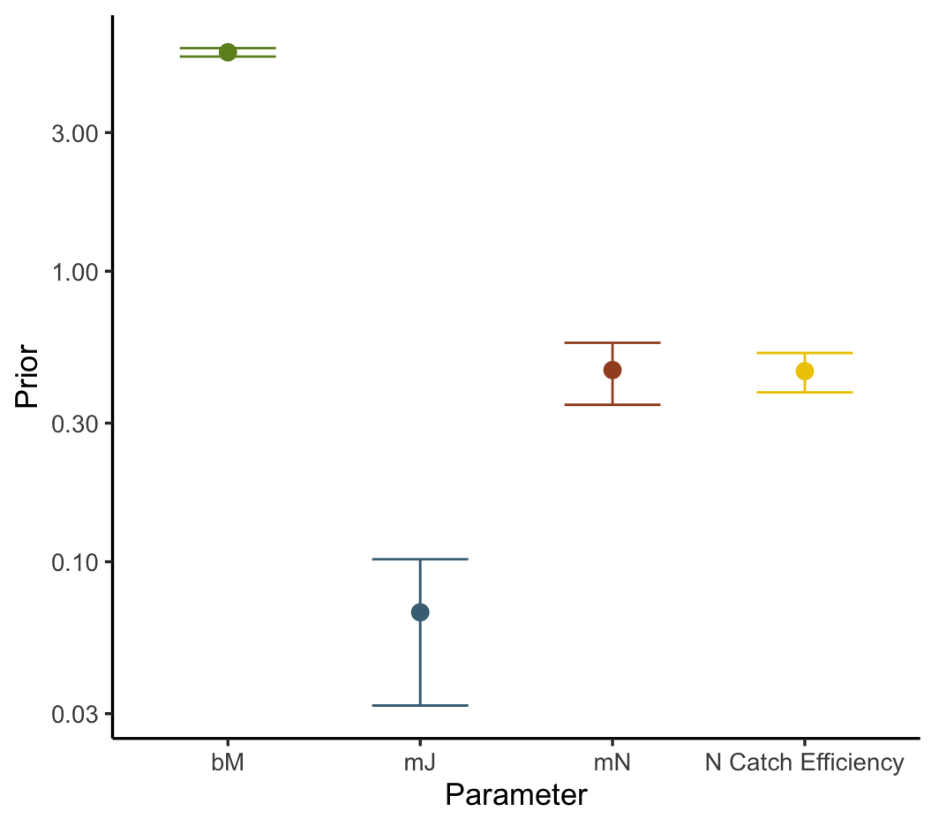

**Figure S2.** Bayesian priors (points and error bars represent means ± SE) for our state space model for copepod birth rates, maturation rates (both nauplii to juvenile and juvenile to adult), and nauplii catch efficiency. The units of birth and maturation rates are per day. Catch Efficiency is a unitless proportion.

**(C) Appendix S3: Parameter estimates**

**Table S1.** Parameters from the chain with the highest posterior density and linear chain trick assumptions to simulate transmission scenarios.

| **Parameter** | **Description** | **Value** | **Source** |
| --- | --- | --- | --- |
| $b_{M}$ | Copepod birthrate | $5.63$ | State space parameter estimates, informed by priors (Fig S2) |
| $d_{A}$ | Adult death rate | $8.04 \times{10}^{-3}$ | State space parameter estimates |
| $m_{J}$ | Maturation rate from juvenile to adult | $1.53 \times{10}^{-1}$ | State space parameter estimates, informed by priors (Fig S2) |
| $d_{J}$ | Relative death rate of juveniles | $1.53 \times{10}^{-3}$ | State space parameter estimates |
| $m_{N}$ | Maturation rate from nauplii to juveniles | $4.1 \times{10}^{-1}$ | State space parameter estimates, informed by priors (Fig S2) |
| $d_{N}$ | Relative death rate of nauplii | $2.15\times{10}^{-1}$ | State space parameter estimates |
| $A_{0}$ | Initial density of adults | $6.39 \times{10}^{2}$ | State space parameter estimates |
| $J_{0}$ | Initial density of juveniles | $1.2 \times{10}^{4}$ | State space parameter estimates |
| $N_{0}$ | Initial density of nauplii | $8.7 \times{10}^{3}$ | State space parameter estimates |
| $\kappa$ | Negative binomial dispersion parameter | $2.64$ | State space parameter estimates |
| ${comp}_{b}$ | Density dependence on births | $3.98 \times{10}^{-6}$ | State space parameter estimates |
| ${comp}_{d}$ | Density dependence on deaths | $8.96 \times{10}^{-4}$ | State space parameter estimates |
| ${comp}_{m}$ | Density dependence on maturation | $3.62 \times{10}^{-3}$ | State space parameter estimates |
| $c_{J}$ | Relative competitive effect of juveniles | $9.94 \times{10}^{-1}$ | State space parameter estimates |
| $c_{N}$ | Relative competitive effect of nauplii | $1.98\times{10}^{-4}$ | State space parameter estimates |
| *cannibalism* | Rate of cannibalism of nauplii by adults | $9.4 \times{10}^{-6}$ | State space parameter estimates |
| *naup_catch* | Nauplii catch efficiency with 100-micron sieve | $4.66 \times{10}^{-1}$ | State space parameter estimates, informed by priors (Fig S2) |
| $l_{s}$ | Number of latent stages between exposure and infection | 60 | Chosen to correspond to latent period estimates(Hurtado and Kirosingh 2019) |
| $l_{R}$ | Rate that an individual copepod moves through each latent stage | 4.3 | Chosen to correspond to latent period estimates(Hurtado and Kirosingh 2019) |
| $\lambda$ | Instantaneous infection rate of adults | 0.032 | Calculated to achieve ~5% infection prevalence amongst susceptibles (non-infected adults)(Lyons 1972, Steib and Mayer 1988, Garrett et al. 2020) |

**(D) Appendix 4: Posterior distributions**

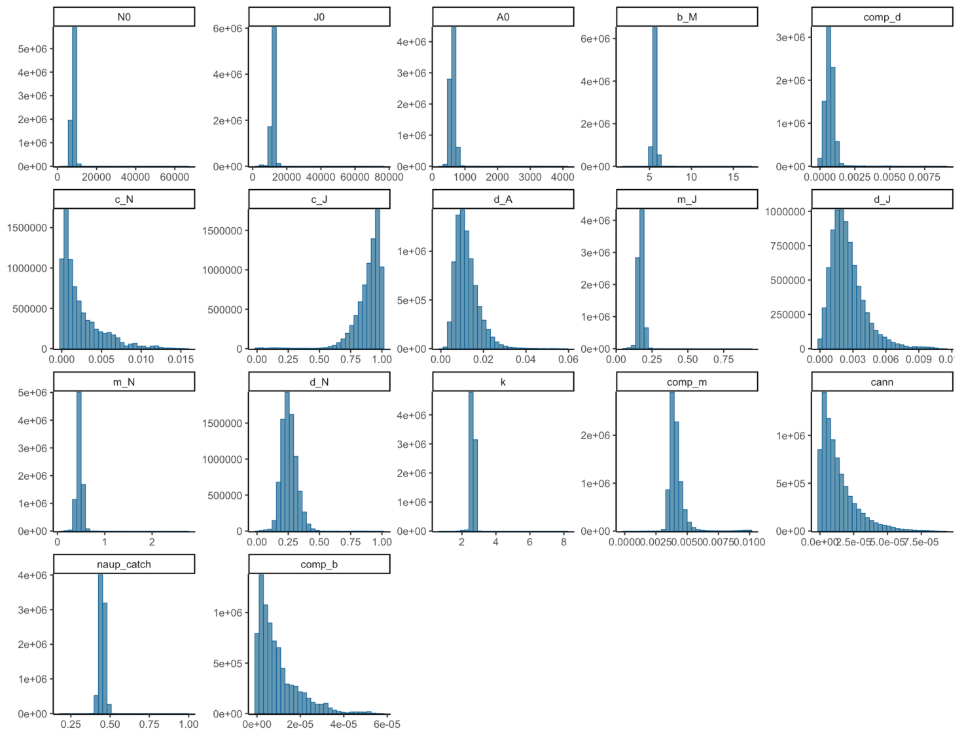

**Figure S1.** Posterior density plots for parameters estimated in the model.

**(E) Appendix 5: Model fit to data**

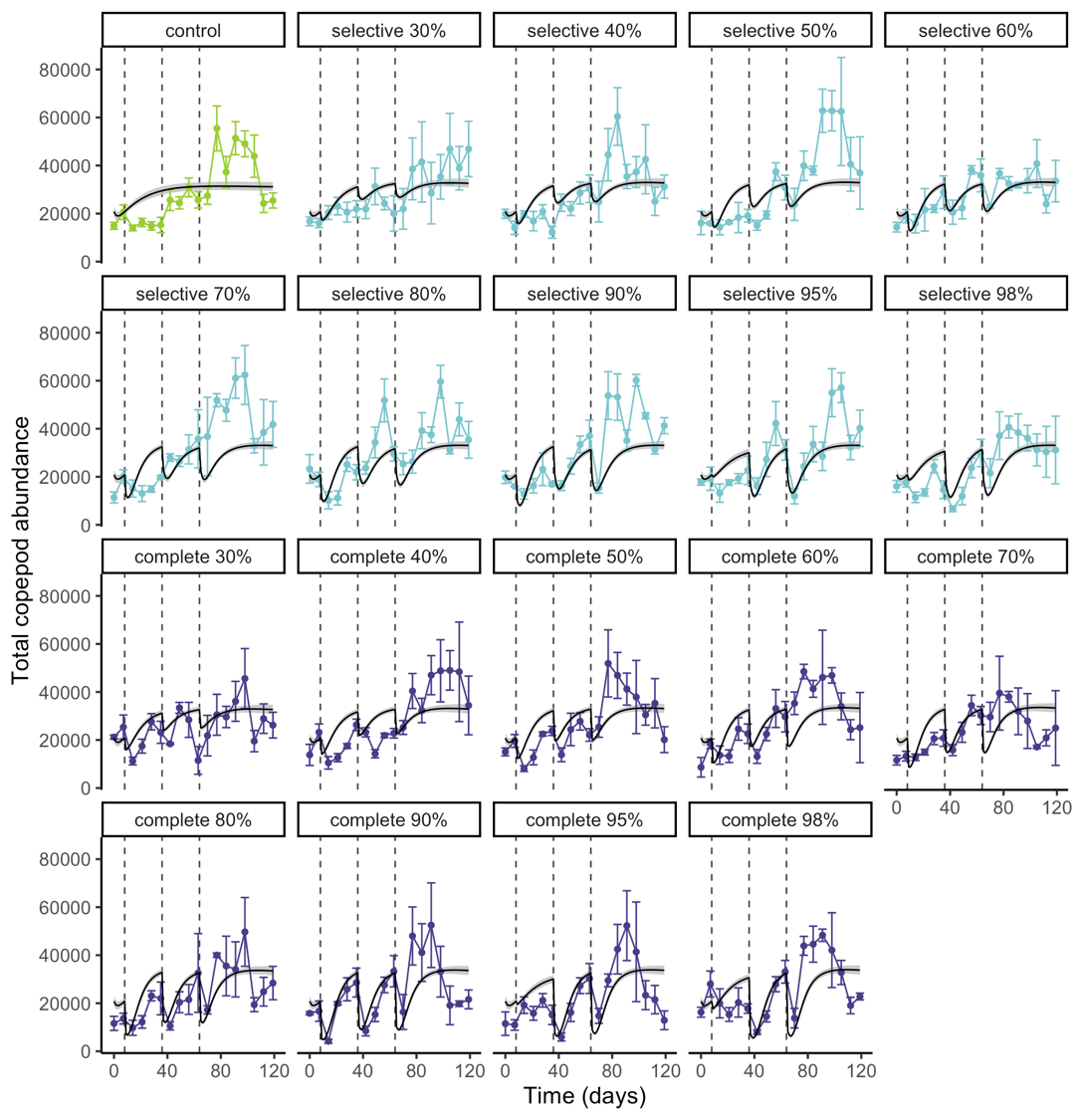

**Figure S1A. Model fit to the data for total copepods.** Lines are treatment means with 95% CI from 1,000 simulations using parameters with maximum log-likelihood. Raw mesocosm data treatment means ± SE plotted in colors.

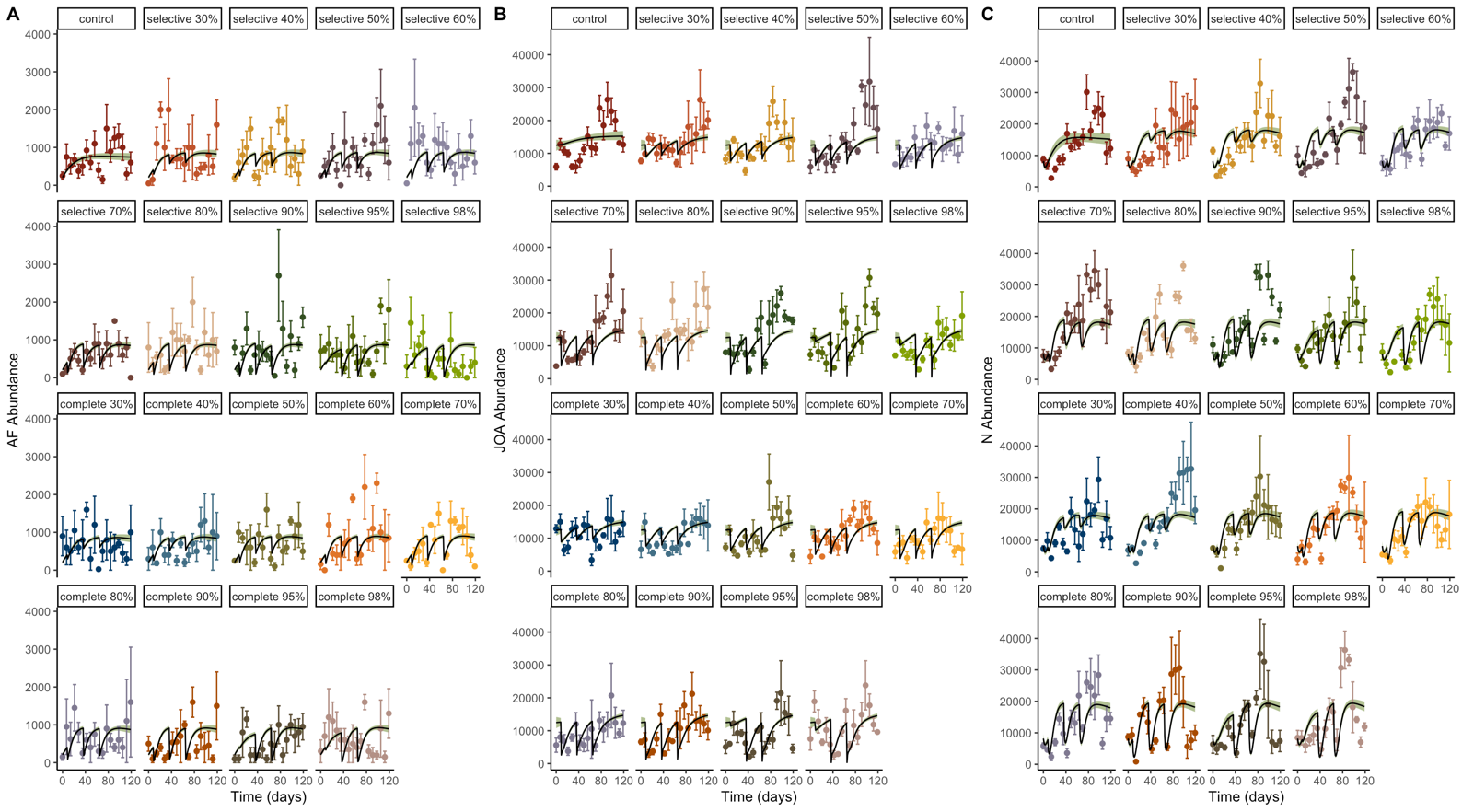

**Figure S1B. Model fit to the data for AF (adult females), JOA (juveniles and other adults), and N (nauplii).** Lines are treatment means with 95% CI from 1,000 simulations using parameters with maximum log-likelihood. Raw mesocosm data treatment means ± SE plotted in colors.

**(F) Appendix 6: Testing for density dependence in reproduction**

**Supplementary Methods S1:** One of our hypothesized mechanisms of density dependence is a reduction in *per capita* birth rate as density increases. We tested for evidence consistent with this mechanism by measuring the females, counting their eggs, and correlating these measures with copepod densities at the time of collection. For each egg-bearing female from each collected sample, we photographed them using a Sedwick rafter slide with 1mm grid under a dissecting microscope at 10-25x and measured individuals using ImageJ. We then stained the copepods with Lugol’s iodine solution and counted eggs by gently pry each egg mass off the copepod and rotating them in all dimensions using a pin. We next wanted to determine if there was a relationship between copepod biomass, total competitive environment, and egg counts. To determine copepod biomass, we used length-weight regression coefficients from Rosen 1981. Specifically, for adults with eggs, eggless individuals, and copepodites, we used values from *Mesocyclops edax* and for nauplii we used Rosen’s estimate for “copepod nauplii”. We measured 10 copepods per stage and used the formulas specified by Rosen to calculate masses. We multiplied the relative abundances of each size class by their masses for each tank over time and summed them up to get total biomass of all copepods for a given tank and time point. To assess the total competitive environment, we multiplied the number of copepods of each life stage per L by the competitive strength of that life stage. With the egg count, biomass, and competitive environment data, we ran two generalized linear mixed models (glmm), one with a fixed effect of total biomass per L and the second with a fixed effect of total competitive environment. Each model used the COM-Poisson distribution with a random effect for tank, week, and copepod id and a response variable of egg count.

**Supplementary Equation S2:**

EggModel = glmmTMB(Egg_Count ~ TotalBiomassPerL + (1 | Tank/Week/Copepod_ID), family=compois, data=Egg_data)

**Supplementary Results S3:** Average calculated masses were 6ug for adults, 1.55 ug for eggless individuals and copepodites, and 0.82 ug for nauplii. We find no significant relationship between egg counts and total biomass or total competitive environment (p=0.446 and 0.192, respectively), consistent with the weak density dependence detected on births in the fitted model.

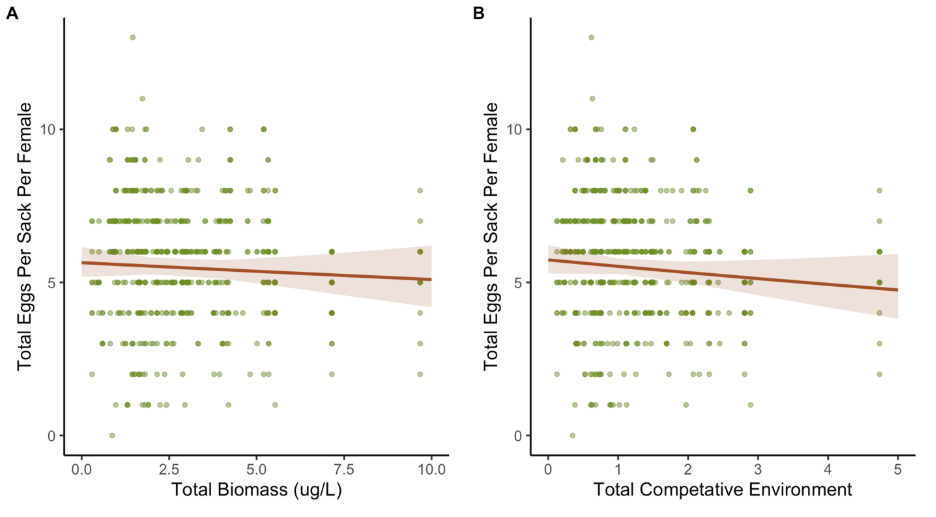

**Supplementary Figure S4. Egg counts for adult females from mesocosm vs. total biomass and total competitive environment.** Green dots denote individual egg masses; red line shows predicted values from GLMM with a 95% confidence band.

**(G) Appendix S7: Evaluating bias and precision of removal copepods removed during interventions.**

**Methods S7.** When we implemented the second and third interventions, experiment days 36 and 64, respectively, we preserved the removed individuals to analyze how well our interventions mapped on to the nominal treatment levels. We anticipated stochasticity involved in removing individuals by removing a portion of habitat, as this is akin to binomial sampling. However, there could also bias in our removal procedures. For example, if copepods could swim away from our collection beaker, then we would remove fewer individuals than anticipated. Thus, for the second and third interventions, we estimated the proportion of individuals removed from a subset of “complete removal” tanks by comparing the abundance of individuals removed to the abundance of individuals estimated the day prior as part of the weekly sampling for the experiment. We then regressed these estimates against the nominal proportions. If the process results in unbiased removal, then the estimated proportions would fall along the 1:1 line with the nominal proportions. Given the similarity of the sample collection, removal, and subsample counting processes to binomial draws, we also simulated the general procedure under basic assumptions of a fixed “true” initial abundance (N_0_ = 16,000 individuals) and fixed quantities for volumes collected (0.5 L for the “pre-intervention” estimate that occurred as part of our weekly sampling and a removal volume corresponding to each treatment, e.g., the 98% removal yields a volume of 14.7 L) and subsample volumes counted that matched the average from the experiment at these time points (1 mL subsample for all “pre-intervention” estimates and 3 mL subsample counted for all “removal” samples). We simulated this process 1000 times for each intervention level (proportions from 0.3 – 0.98 removal) and then calculated the mean and 95% confidence intervals for the expected range of values.

We first concentrated all removed individuals into a 50-mL centrifuge tube with 92.5% EtOH. Because these removal volumes could be as large as 14.7 L, we diluted these samples to 50 mL, rather than 10 mL. We then moved a 10 mL sample to a new tube and subsample to a new tube. We then counted 1-mL at a time until 50 total individuals were counted.

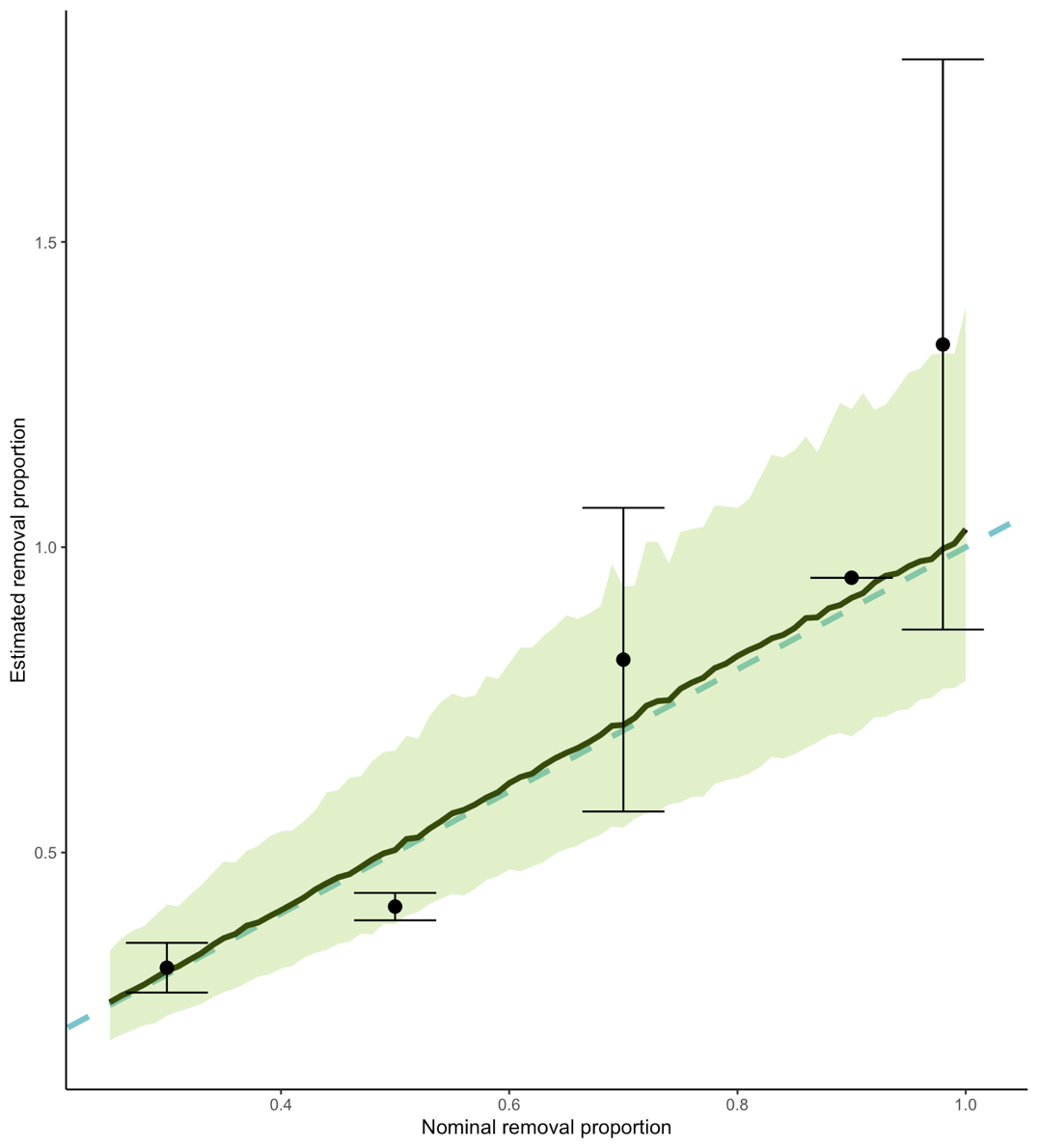

**Figure S1.** Treatment means and SE across interventions two and three plotted against means and 95% CI for simulated data across the removal gradient. The dashed light blue line is the 1:1 line. We generally found agreement between the nominal and calculated estimates. Any slight disagreement reflects the stochasticity of sampling. Results shown for the complete removal samples.

**Literature Cited**

Garrett, K. B., E. K. Box, C. A. Cleveland, A. A. Majewska, and M. J. Yabsley. 2020. Dogs and the classic route of Guinea Worm transmission: an evaluation of copepod ingestion. Sci Rep **10**:1430.

Hurtado, P. J., and A. S. Kirosingh. 2019. Generalizations of the 'Linear Chain Trick': incorporating more flexible dwell time distributions into mean field ODE models. J Math Biol **79**:1831-1883.

Lyons, G. R. 1972. Guineaworm infection in the Wa district of north-western Ghana. Bull World Health Organ **47**:601-610.

Steib, K., and P. Mayer. 1988. Epidemiology and vectors of Dracunculus medinensis in northwest Burkina Faso, West Africa. Annals of Tropical Medicine & Parasitology **82**:189-199.
